# Acid ceramidase inhibition enhances BCL-2 targeting in venetoclax-resistant acute myeloid leukemia via a cytotoxic integrated stress response

**DOI:** 10.1101/2025.06.06.657881

**Authors:** Johnson Ung, Su-Fern Tan, Jeremy J.P. Shaw, Maansi Taori, Tess M. Deddens, Giovana da Costa Venancio, McLane M. Montgomery, James T. Hagen, Raphael T. Aruleba, Upendar R. Golla, Arati Sharma, B. Bishal Paudel, Irene Sung-Ah Lee, Bhavishya Ramamoorthy, Kevin A. Janes, Francine Garrett-Bakelman, Myles C. Cabot, Kelsey H. Fisher-Wellman, Todd E. Fox, David F. Claxton, Charles E. Chalfant, David J. Feith, Thomas P. Loughran

## Abstract

Resistance to combination regimens containing the BCL-2 inhibitor venetoclax in acute myeloid leukemia (AML) is a growing clinical challenge for this extensively utilized agent. We previously established the anti-leukemic properties of ceramide, a tumor-suppressive sphingolipid, in AML and demonstrated that upregulated expression of acid ceramidase (AC), a ceramide-neutralizing enzyme, supported leukemic survival and resistance to BH3 mimetics. Here, we report the anti-leukemic efficacy and mechanisms of co-targeting AC and BCL-2 in venetoclax-resistant AML. Analysis of the BeatAML dataset revealed a positive relationship between increased AC gene expression and venetoclax resistance. Targeting AC enhanced single-agent venetoclax cytotoxicity and the venetoclax + cytarabine combination in AML cell lines with primary or acquired venetoclax resistance. SACLAC + venetoclax was equipotent to the combination of venetoclax + cytarabine at reducing cell viability when evaluated *ex vivo* across a cohort of 71 primary AML patient samples. Mechanistically, SACLAC + venetoclax increased ceramide to levels that trigger a cytotoxic integrated stress response (ISR), ISR-mediated NOXA protein upregulation, mitochondrial dysregulation, and caspase-dependent cell death. Collectively, these data demonstrate the efficacy of co-targeting AC and BCL-2 in AML and rationalize targeting AC as a therapeutic approach to overcome venetoclax resistance.

## Introduction

Acute myeloid leukemia (AML) is a devastating hematologic malignancy with limited treatment options and poor survival outcomes (1). Dysregulated mitochondrial apoptosis supports AML resistance to anti-neoplastic agents (2). Intracellular cell death signals converge in part at the mitochondria where cell death is regulated by the coordinated activity of anti-apoptotic (e.g., BCL-2, MCL-1, BCL-xL) and pro-apoptotic (e.g., BIM, NOXA, BAX, BAK) BCL-2 family proteins (3). A targetable BCL-2 vulnerability was identified in AML and spurred clinical trials evaluating the BCL-2 inhibitor venetoclax alone or in multiple combinations (4). Modestly active as a monotherapy (5), venetoclax was effective in combination with hypomethylating agents or low-dose cytarabine in early studies, which supported its accelerated approval in 2018 (6, 7). Positive results from the confirmatory phase 3 VIALE-A and VIALE-C clinical trials evaluating venetoclax + azacitidine versus azacitidine alone or venetoclax + low-dose cytarabine versus cytarabine alone supported the regular approval of venetoclax in 2020 (8, 9). The adoption of venetoclax-containing regimens to intensive combination chemotherapy was especially beneficial for chemotherapy-ineligible AML patients. Unfortunately, responses to venetoclax-containing regimens are transient and resistance is an unmet clinical challenge (10). Therefore, identifying additional vulnerabilities to enhance venetoclax-containing regimens is paramount.

Sphingolipids are a heterogeneous class of structural and signaling lipids (11). Important for oncology is the sphingolipid species ceramide, the tumor-suppressive properties of which were initially described in myeloid leukemia as a potential therapeutic vulnerability (12). Ceramides reside at the canonical center of sphingolipid metabolism and are generated via *de novo* synthesis, higher-order sphingolipid catabolism, or inhibition of ceramide-catabolizing pathways (13). Ceramides contribute to the cytotoxic mechanism of AML therapeutics including cytarabine (14), daunorubicin (15, 16), BH3 mimetics (17), and FLT3 inhibitors (18). We previously demonstrated that exogenous ceramide supplementation through ceramide nanoliposomes augmented the efficacy of venetoclax and cytarabine in combination (19). Heightened ceramide depletion is an emerging AML feature due in part to upregulation of acid ceramidase (AC), a lysosomal lipid hydrolase and ceramide-catabolizing enzyme (20). Previous work in AML from our group found that AC upregulation supports leukemic survival and resistance to chemotherapy and BH3 mimetics (17, 21). While genetic and pharmacologic AC inhibition resensitized drug-resistant AML cells to chemotherapy, the effect of AC inhibition on BH3 mimetic efficacy and venetoclax-containing regimens remained unknown.

In this study, we evaluated AC and BCL-2 co-targeting using cell death assays, western blotting, sphingolipid profiling, and rescue studies to define the efficacy and cytotoxic mechanisms. Co-targeting AC and BCL-2 induced ceramide accumulation and a cytotoxic integrated stress response leading to mitochondrial dysfunction and caspase-dependent cell death. Collectively, this work highlights the importance of ceramide clearance in modulating venetoclax cytotoxicity and provides the rationale for pursuing AC-targeting modalities to augment venetoclax-containing regimens.

## Materials and Methods

### Cell culture

Cells were cultured in RPMI-1640 (Corning, NY, USA, #10-040; for MM-6) or IMDM (ThermoFisher, Waltham, MA, USA, #12440; for MV4-11 variants) supplemented with 10% FBS (VWR, Radnor, PA, USA, #97068–085) and incubated in humified incubators set to 37°C with 5% CO_2_. MM-6 cells were purchased from DSMZ. HL-60 and MV4-11 cells were obtained from ATCC (Manassas, VA, USA). Venetoclax-resistant MV4-11 cells (MV4-11 VEN-R) were generated by our group via continued subculture in media containing venetoclax at 0.25 µM. HL-60 cells were transduced to express GFP and luciferase as described in the supplemental methods. MV4-11 cells expressing YFP and luciferase were kindly gifted by Kenichiro Doi (22). Primary AML patient sample acquisition, processing, and handling are detailed in the supplement.

### Cell viability assays

For cell lines: cells were seeded in 96-well flat-bottom plates at 2×10^5^ cells/mL and treated at the indicated concentrations in a final volume of 100 μL. Following treatment, 20 μL of [3-(4,5-dimethylthiazol-2-yl)-5-(3-carboxymethoxyphenyl)-2-(4-sulfophenyl)-2H-tetrazolium] (MTS) (Promega, Madison, WI, USA, #G3582) was added and incubated for an additional 3 h. Absorbance was measured at 490 nm with the BioTek Cytation 3 plate reader (Biotek, Winooski, VT, USA). Following background subtraction, responses were normalized to vehicle control which was defined as 100%. For patient samples: Single and combination drug screening for patient samples was performed as previously described (23). Briefly, 5000 cells were seeded in 384-well plates and treated at the indicated concentrations for the indicated times. Cell viability was determined using the CellTiter-Glo luminescence assay (Promega, #G755B), and luminescence was measured using GloMax Discover (Promega).

### Flow cytometry

Cells were seeded at 2.5×10^5^-5×10^5^ cells/mL and treated as indicated in 6-well plates or tissue culture flasks. Following treatment, cells were assayed per the manufacturer’s instructions using the Muse Annexin V & Dead Cell Kit (Cytek, Fremont, CA, USA, #MCH100105), Muse MitoPotential Kit (Cytek, #MCH100110), or Muse Count & Viability Kit (Cytek, #MCH100102) per the manufacturer’s protocol. For all assays, cells were diluted to <5×10^5^ cells/mL and 1000-2000 events were captured using the Cytek Guava Muse Cytometer. Data were analyzed using the Muse software version 1.8.0.3. Each assay identified live, apoptotic, and dead cells.

### Western blotting

Cells were seeded at 2.5×10^5^-5×10^5^ cells/mL and treated as indicated in 6-well plates or tissue culture flasks. Following treatment, cells were centrifuged at 400xg for 7 min, washed in 1X PBS, and subjected to protein isolation and western blotting as previously described (23).

### Cytotoxicity rescue studies

Cells were seeded at 2.5×10^5^-5×10^5^ cells/mL and pretreated as indicated in 6-well plates or tissue culture flasks. Following pretreatment, cells were treated with vehicle, SACLAC, venetoclax, or the combination at the indicated concentrations and times and assayed.

### Sphingolipid profiling

Cells were seeded at 2.5×10^5^-5×10^5^ cells/mL and treated as indicated in 6-well plates or tissue culture flasks. Following treatment, sphingolipids were extracted, processed, and quantified by LC-MS as previously described (23).

### Quantification and statistical analysis

Statistical tests were performed as indicated in figure legends with GraphPad Prism (version 10.4.2). Dose response curves were generated and analyzed using the Prism function “log(inhibitor) vs. response—variable slope (four parameters)”. Bliss synergy analyses were performed using SynergyFinder 2.0 as previously described (23). Experiments were performed in three independent replicates with three or more technical replicates. Unless otherwise stated, a representative experiment is shown with error bars representing mean ± standard deviation (SD).

## Results

We previously established a relationship between AC overexpression and resistance to the BH3 mimetic ABT-737 (17). To explore links between AC and the clinically approved BH3 mimetic venetoclax, we analyzed the BeatAML database (24) and identified a positive association between increased AC mRNA expression (gene name *ASAH1*) and increased venetoclax resistance (**Fig. 1a**). These findings prompted us to study the therapeutic potential of co-targeting AC and BCL-2 in AML cell lines and primary AML patient samples using two previously characterized small molecule AC inhibitors: SACLAC (22) and LCL-805 (23)).

**Fig. 1:**
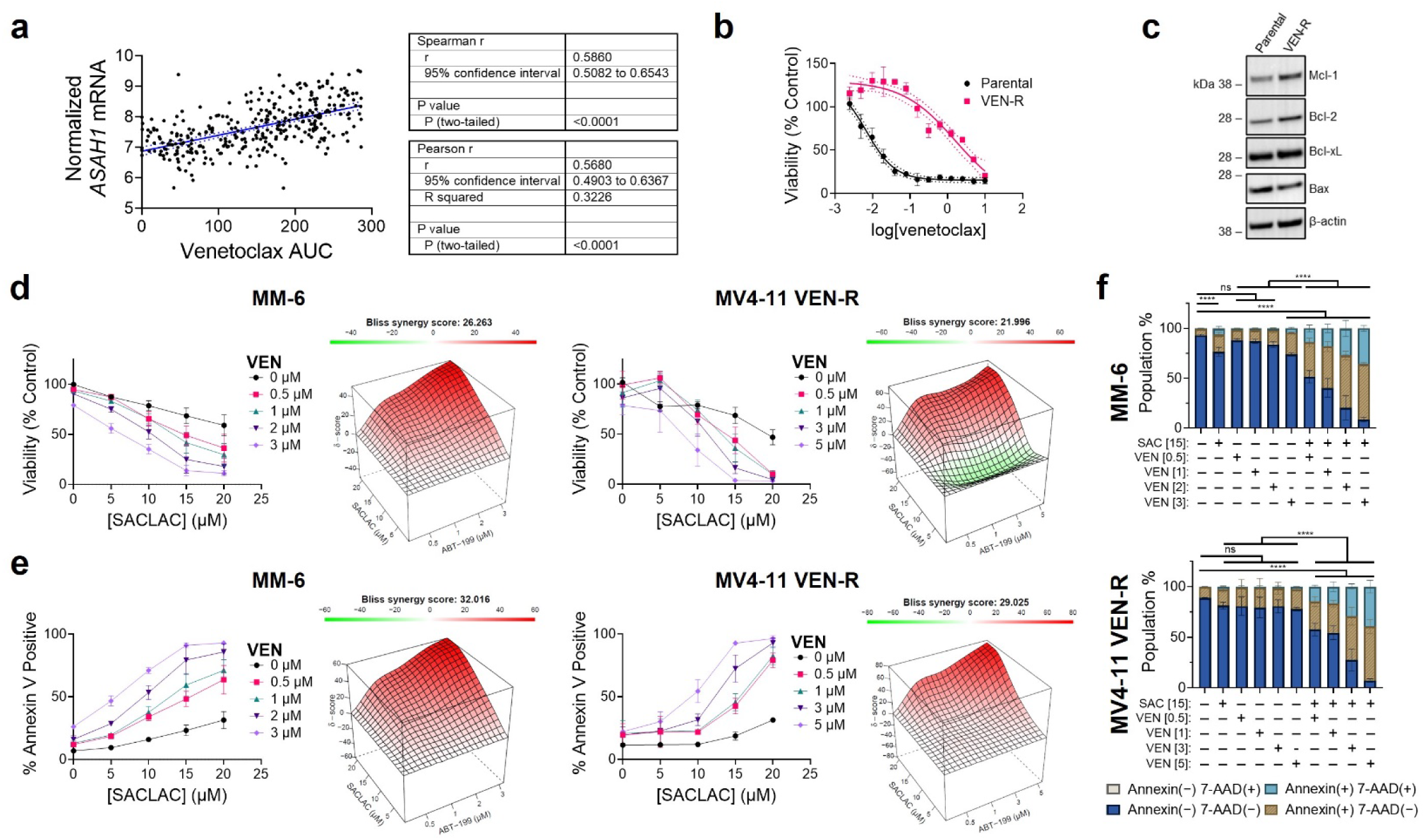
Pharmacologic inhibition of acid ceramidase enhanced venetoclax cytotoxicity in venetoclax-resistant acute myeloid leukemia cell lines. **A** Correlation analyses comparing venetoclax area under the curve (AUC) values and *ASAH1* mRNA expression (n=332) from BeatAML2.0 (http://www.vizome.org/aml2/). Dotted lines represent linear regression with 95% confidence interval. **B** MV4-11 Parental and MV4-11 VEN-R cells were subjected to increasing concentrations of venetoclax for 48 h and cell viability was assessed by MTS. Dotted lines represent 95% confidence interval. **C** Immunoblotting of MV4-11 Parental and MV4-11 VEN-R cells. β-actin expression served as the loading control for immunoblots. **D, E** MTS cell viability, flow cytometry annexin V staining, and Bliss synergy scores (SynergyFinder 2.0) were measured following DMSO, SACLAC, venetoclax, or combination treatment at the indicated concentrations in MM-6 (48 h) and MV4-11 VEN-R cells (72 h). Bliss scores greater than 10 were considered synergistic, between −10 and 10 were considered additive, and less than −10 were considered antagonistic. **F** MM-6 and MV4-11 VEN-R cells were treated with DMSO, SACLAC, venetoclax, or the combination for 48 h or 72 h, respectively, and annexin V and 7-AAD population percentages were measured by flow cytometry. Significance was assessed by two-way ANOVA with Tukey’s multiple comparisons test comparing live cells (annexin− 7-AAD−) across treatment groups. Data are presented as mean ± SD of three independent experiments. ABT-199 = venetoclax (VEN). Concentrations in brackets are µM. ****p<0.0001, ns = not significant.

### AC inhibition enhanced BCL-2 sensitivity in venetoclax-resistant AML cells

Primary and acquired mechanisms contribute to venetoclax resistance (25). To model both, we evaluated the efficacy of co-targeting AC and BCL-2 in human AML cell lines with primary (MM-6) or acquired venetoclax resistance (MV4-11 VEN-R; generated through continual venetoclax exposure). Compared to parental MV4-11 cells, MV4-11 VEN-R cells were venetoclax resistant and exhibited increased protein levels of the anti-apoptotic proteins MCL-1, BCL-2, and BCL-xL as well as decreased levels of the pro-apoptotic protein BAX (**Fig. 1b-c**). To test whether the AC targeting enhances venetoclax cytotoxicity, we exposed AML cells to SACLAC and venetoclax at increasing concentrations and assessed cell viability and annexin V/7-AAD staining. SACLAC and venetoclax significantly reduced cell viability (**Fig. 1d**) and increased the fraction of annexin V-positive cells (**Fig. 1e**) in a concentration-dependent manner in MM-6 and MV4-11 VEN-R cells. Combination effects were quantified with SynergyFinder 2.0 (26) and yielded Bliss scores indicative of strong synergy (**Fig. 1d-e**). Additional flow cytometry profiling of SACLAC and venetoclax-treated cells demonstrated significantly reduced live cells and increased annexin V positive, 7-AAD positive, and double-positive cells (**Fig. 1f**). Co-targeting AC and BCL-2 also enhanced cell killing in venetoclax-sensitive human AML cell lines (**Fig. S1**). Together, these findings show that AC inhibition significantly enhances venetoclax cytotoxicity in venetoclax-sensitive and - resistant AML cells.

### Co-targeting AC and BCL-2 resulted in ceramide accumulation, caspase activation, and NOXA protein upregulation

We extended our studies to investigate the cytotoxic mechanism underlying AC and BCL-2 co-targeting. Ceramide accumulation following inhibition of anti-apoptotic BCL-2 family proteins potentiates apoptotic cell death (27) and ceramide nanoliposomes exacerbated venetoclax cytotoxicity in AML (19). Thus, we next assessed the impact of AC and BCL-2 inhibition on intracellular ceramide levels. In both MM-6 and MV4-11 VEN-R, SACLAC treatment increased total intracellular ceramide levels by 4-fold while venetoclax alone increased levels by 1.5-fold. Ceramides of C16 and C24:1 fatty acid chain length were the primary species increased. Combining SACLAC and venetoclax resulted in significantly greater ceramide accumulation over vehicle (MM-6=7.3-fold; MV4-11 VEN-R=3.7-fold), SACLAC (MM-6=1.3-fold; MV4-11 VEN-R=1.5-fold), or venetoclax (MM-6=3.8-fold; MV4-11 VEN-R=2.8-fold) (**Fig. 2a**). SACLAC treatment elevated hexosylceramide levels in MV4-11 VEN-R cells while the drug combination reduced sphingomyelin levels in MM-6 cells (**Fig. S2)**. These findings demonstrate that AC inhibition and venetoclax treatment cooperate to elevate intracellular ceramide accumulation to drive cytotoxicity.

**Fig. 2:**
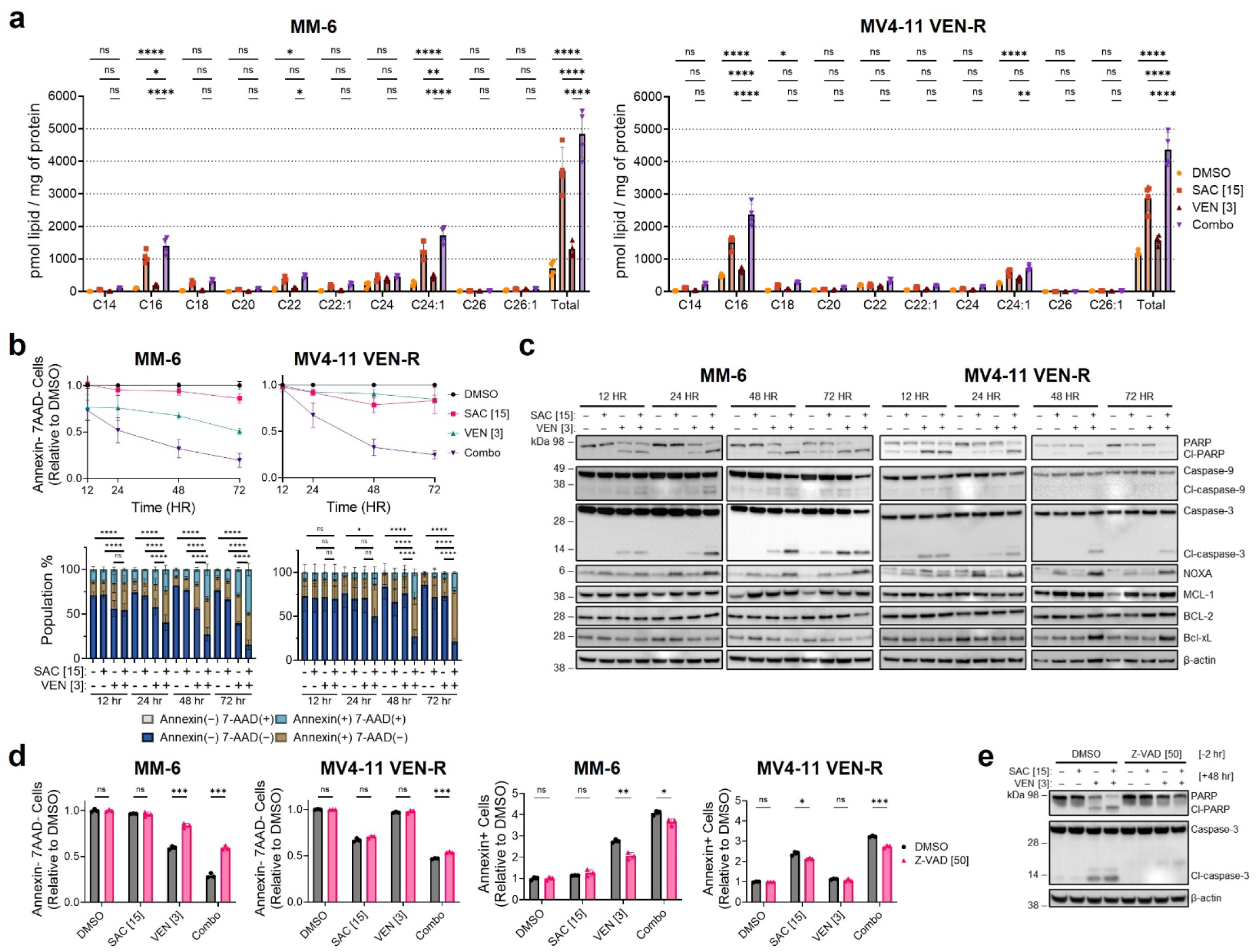
SACLAC and venetoclax induced ceramide accumulation and caspase-mediated cell death characterized by NOXA accumulation. **A** MM-6 and MV4-11 VEN-R cells were treated with DMSO, SACLAC, venetoclax, or the combination, and ceramide levels were measured by liquid chromatography-mass spectrometry. Significance was assessed by two-way ANOVA with Tukey’s multiple comparisons test. **B, C** Flow cytometry annexin V and 7-AAD profiling and immunoblotting of MM-6 and MV4-11 VEN-R cells treated with DMSO, SACLAC, venetoclax, or the combination at the indicated concentrations for 12, 24, 48, and 72 h. Quantitative data are the mean of three independent experiments. Significance was assessed by two-way ANOVA with Tukey’s multiple comparisons test comparing live cells (annexin− 7-AAD−) across treatment groups. **D** Flow cytometry annexin V profiling of cells pretreated with DMSO or Z-VAD-FMK for 2 h followed by DMSO, SACLAC, venetoclax, or the combination treatment for 48 h. Significance was assessed by unpaired t-test with Welch’s correction. **E** Immunoblotting of MM-6 cells pretreated with DMSO or Z-VAD-FMK for 2 h followed by DMSO, SACLAC, venetoclax, or the combination treatment for 48 h. β-actin expression served as the loading control for immunoblots. Data are presented as mean ± SD. SAC = SACLAC; VEN = venetoclax; Z-VAD = Z-VAD-FMK. Concentrations in brackets are µM. *p<0.05, **p<0.01, ***p<0.005, ****p<0.0001, ns = not significant.

We next determined the effects of SACLAC and venetoclax treatment on apoptotic and BCL-2 family protein levels over a time course. SACLAC enhanced venetoclax cytotoxicity in a time-dependent manner across both cell lines beginning 24 h post treatment as measured by reduced live cell numbers and enhanced annexin V, positive, 7-AAD, and double positive cell percentages (**Fig. 2b**). Cell death following AC and BCL-2 inhibition was accompanied by increased PARP and caspase-3 cleavage, suggestive of caspase activation (**Fig. 2c**). To test whether caspase activation was necessary to induce cell death with SACLAC + venetoclax, we pretreated cells with the pan-caspase inhibitor Z-VAD-FMK followed SACLAC, venetoclax, or the combination. Z-VAD-FMK pretreatment protected against venetoclax and SACLAC + venetoclax cell killing in MM-6 cells concomitant with reduced PARP and caspase-3 cleavage (**Fig. 2d-e**). However, Z-VAD-FMK conferred less protection to SACLAC + venetoclax in MV4-11 VEN-R cells. Despite increased cell death, anti-apoptotic BCL-2 family protein levels were not diminished. Instead, we observed increased protein levels of the pro-apoptotic BH3-only protein, NOXA, a well-described endogenous MCL-1 inhibitor (**Fig. 2c**). These results demonstrate that co-targeting AC and BCL-2 enhanced caspase-mediated cell death and upregulated the BH3-only protein NOXA.

### AC and BCL-2 co-targeting induced a cytotoxic integrated stress response

To better understand how SACLAC modulates venetoclax cytotoxicity, we examined the proteome following SACLAC treatment in MV4-11 cells and identified 249 differentially upregulated and 190 differentially downregulated proteins (**Fig. 3a**). STRINGDb pathway analysis identified proteins involved in “translation”, “ribonucleoprotein complex biogenesis”, and “ribonucleoprotein complex assembly” Gene Ontology term processes as significantly depleted following SACLAC treatment (**Fig. 3b, Table S1**). These data are consistent with previous reports that ceramides induce the integrated stress response (ISR), which regulates translation in response to cellular stress (28). ISR activation also sensitizes AML cell lines to venetoclax (28, 29). To evaluate the link between the ISR and co-targeting AC and BCL-2, we measured ISR activation following SACLAC and venetoclax treatment. Central to ISR activation is the phosphorylation of eIF2α at Ser51 (p-eIF2α) by PKR, PERK, GCN2, or HRI, which results in reduced global protein synthesis and upregulation of the transcription factor ATF4 (30). Both SACLAC and venetoclax increased p-eIF2α (Ser51) and ATF4 protein expression in MM-6 and MV4-11 VEN-R cells (**Fig. 3c**). Interestingly, SACLAC + venetoclax treatment cooperated to induce a heightened ISR compared to either single agents. Paradoxically, ISR activation has been reported to support (31, 32) and overcome (28, 33, 34) therapy resistance in AML. To determine whether ISR activation was cytoprotective or cytotoxic, we treated with the small molecule ISR inhibitor ISRIB and assessed ISR signaling and cytotoxicity with subsequent SACLAC and venetoclax treatment. ISRIB pretreatment significantly abrogated SACLAC + venetoclax-mediated cytotoxicity and reduced ATF4 protein upregulation (**Fig. 3d-e**). We validated our findings with a separate AC inhibitor, LCL-805, which also enhanced venetoclax-mediated cell killing in a caspase- and ISR-dependent manner (**Fig. S3a-c**). These findings demonstrate that co-targeting AC and BCL-2 induces a cytotoxic ISR.

**Fig. 3:**
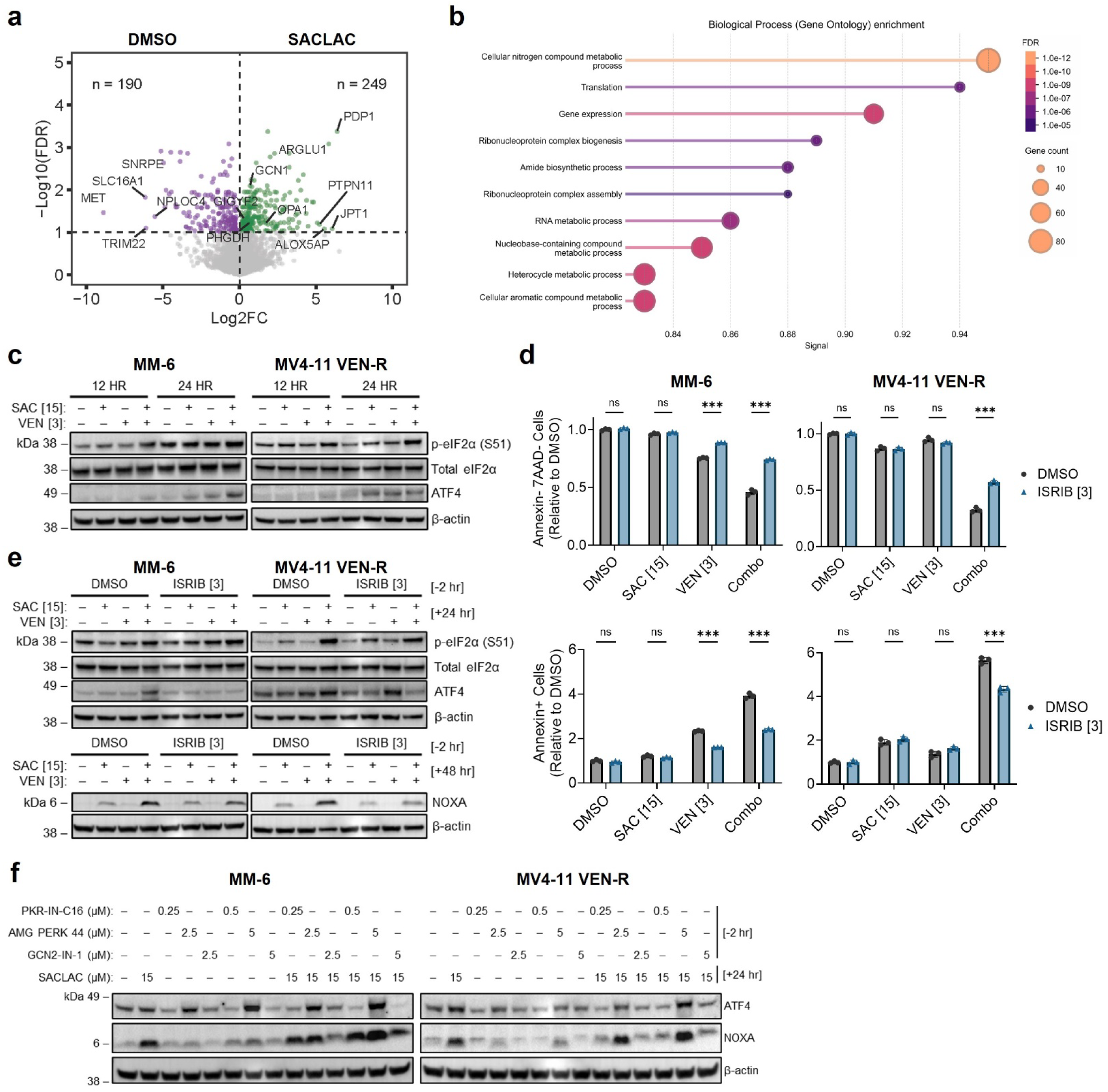
SACLAC and venetoclax upregulated NOXA via a cytotoxic integrated stress response. **A** Volcano plot of significantly enriched (n=249) or depleted (n=190) proteins from SACLAC versus DMSO-treated MV4-11 cells. Significance was defined using an FDR (q)<0.1 cutoff. **B** Biological Process (Gene Ontology) analysis of significantly downregulated proteins from panel A. **C** Immunoblotting of cells treated with DMSO, SACLAC, venetoclax, or the combination for 12 or 24 h. **D** Flow cytometry annexin V profiling of cells pretreated with DMSO or ISRIB for 2 h followed by DMSO, SACLAC, venetoclax, or the combination treatment for 48 h. Significance was assessed by unpaired t-test with Welch’s correction. **E** Immunoblotting of cells pretreated with DMSO or ISRIB for 2 h followed by DMSO, SACLAC, venetoclax, or combination treatment for 24 or 48 h. **F** Immunoblotting of cells pretreated with DMSO, PKR-IN-C16 (PKR inhibitor), AMG PERK 44 (PERK inhibitor), or GCN2-IN-1 (GCN2 inhibitor) for 2 h followed by DMSO or SACLAC treatment for 24 h. β-actin expression served as the loading control for immunoblots. Data are presented as mean ± SD. SAC = SACLAC; VEN = venetoclax. Concentrations in brackets are µM. *p<0.05, ***p<0.005, ns = not significant.

### Integrated stress response activation promoted NOXA protein upregulation

We next studied the relationship between AC inhibition, the ISR, and BCL-2 family proteins to understand the mechanisms underlying AC and BCL-2 co-targeting. Mitochondrial outer membrane permeabilization and apoptosis are regulated by pro-apoptotic and anti-apoptotic BCL-2 proteins (3). Anti-apoptotic proteins were not decreased following treatment with SACLAC or the SACLAC + venetoclax combination. Instead, we observed upregulation of the pro-apoptotic BH3-only protein NOXA (**Fig. 2c**). ISR activation has previously been reported to induce NOXA induction (28, 34, 35), which prompted us to evaluate this link in our model. To test this, we pretreated MM-6 and MV4-11 VEN-R cells with ISRIB and measured NOXA protein following SACLAC and venetoclax treatment. ISRIB pretreatment partially prevented NOXA accumulation following SACLAC + venetoclax treatment compared to cells pretreated with vehicle (**Fig. 3e**). Mechanistically, ISRIB functions downstream of eIF2α phosphorylation by titrating the formation of eIF2B heterodimers to aid eIF2•GTP•methionyl-initiator tRNA ternary complex formation to support translation (30). As an orthogonal approach, we pharmacologically inhibited ISR sensor kinases to determine which pathway was involved in ISR activation following AC inhibition. While the PERK inhibitor, AMG PERK 44, did not blunt SACLAC-induced ATF4 or NOXA protein upregulation, GCN2 inhibition with GCN2-IN-1 significantly reduced ATF4 and NOXA protein upregulation in both cell lines (**Fig. 3f**). PKR inhibition with PKR-IN-C16 also prevented ATF4 accumulation, though the effect on NOXA was only observed in MV4-11 VEN-R cells. Consistent with GCN2 involvement in SACLAC-induced ISR, our proteomics data demonstrated significant enrichment of GCN1, an endogenous GCN2 activator involved in the detection of colliding ribosomes (36) and GIGYF2, which is involved in the ribosome-associated quality control pathway to regulate proteostasis (37) (**Fig. 3a**). Together, these results are consistent with AC inhibition inducing a GCN2- and PKR-mediated ISR leading to NOXA protein upregulation.

### Co-targeting AC and BCL-2 antagonized mitochondrial function

Mitochondrial reprogramming is a hallmark of venetoclax resistance (38) and targeting mitochondrial translation (33) or structure (39) enhances venetoclax cytotoxicity. Because AC inhibition also antagonizes mitochondrial function (40), we posited that AC and BCL-2 inhibition converge at the mitochondria as a combined cytotoxic mechanism. To test this, we assessed the effect of co-targeting AC and BCL-2 on mitochondrial membrane potential, a biomarker of mitochondrial health. While transient depolarization can be tolerated, long-term depolarization commits cells to apoptotic cell death (41, 42). We treated cells with SACLAC and venetoclax and assessed mitochondrial polarization with tetramethylrhodamine, methyl ester (TMRM), a fluorescent dye that localizes to polarized mitochondria. We observed significant mitochondrial depolarization following SACLAC + venetoclax treatment compared to single agent controls in both MM-6 and MV4-11 VEN-R cells (**Fig. 4a**). To determine whether the observed mitochondrial depolarization preceded cell death, we pretreated cells with vehicle or Z-VAD-FMK, treated cells with SACLAC and venetoclax, and assessed mitochondrial polarization. Although Z-VAD-FMK protected against SACLAC + venetoclax-induced cytotoxicity (**Fig. 2d**), Z-VAD-FMK pretreatment did not affect mitochondrial depolarization suggesting that depolarization was upstream of cell death (**Fig. 4a**). As ISR inhibition protected against SACLAC + venetoclax cytotoxicity (**Fig. 3b**), we also tested whether ISRIB affected mitochondrial depolarization induced by the combination. ISRIB pretreatment significantly protected against mitochondrial depolarization induced by SACLAC + venetoclax in MM-6 and MV4-11 VEN-R cells (**Fig. 4b**). These findings suggest that the cytotoxic mechanism of SACLAC + venetoclax converges at and leads to mitochondrial depolarization, enhanced cell death via the ISR, and downstream caspase activation.

**Fig. 4:**
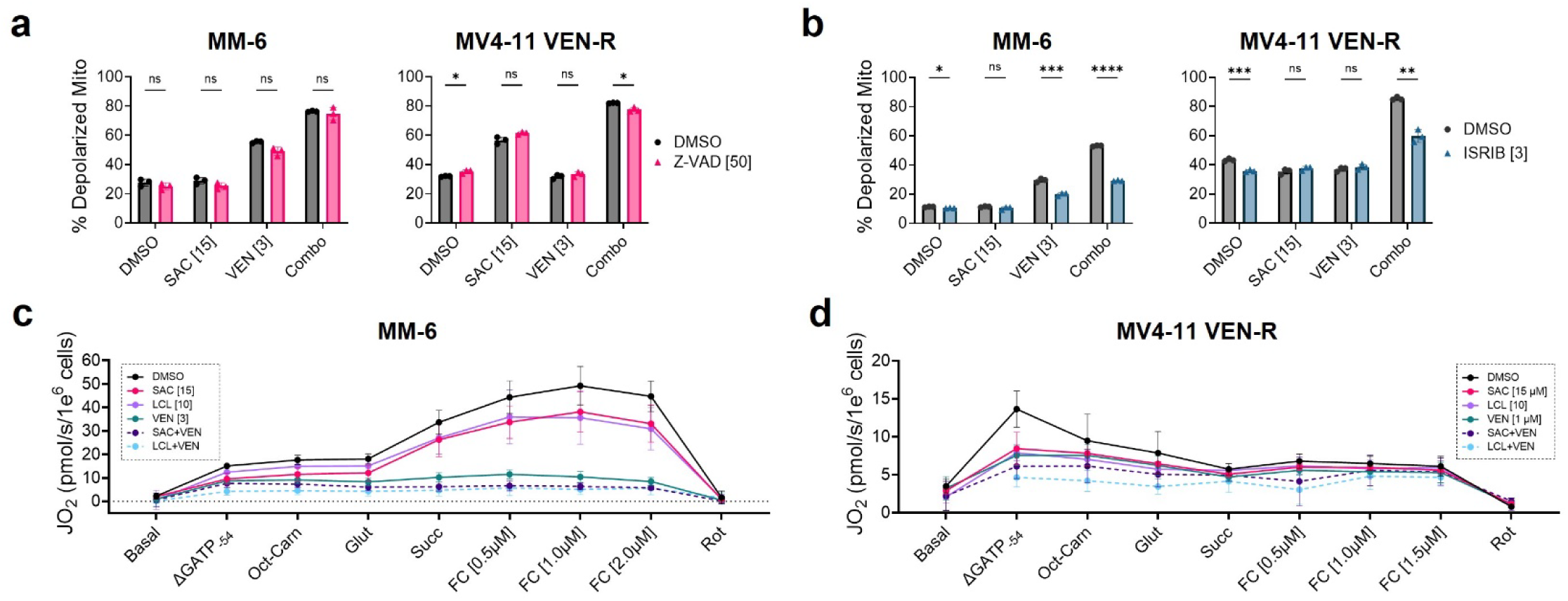
Effect of AC and BCL-2 inhibition on mitochondrial function and membrane potential. **A, B** MM-6 and MV4-11 VEN-R cells were pretreated with DMSO, Z-VAD-FMK, or ISRIB for 2 h then treated with DMSO, SACLAC, venetoclax, or the combination for 48 h. Mitochondrial membrane potential was assessed by flow cytometry. Significance was assessed by unpaired t-test with Welch’s correction. **C, D** MM-6 and MV4-11 VEN-R cells were treated with DMSO, SACLAC, LCL-805, venetoclax, SACLAC + venetoclax, or LCL-805 + venetoclax. Oxygen consumption was measured in digitonin (0.02 mg/mL) permeabilized cells in the absence of substrates (basal), with ATP free energy clamped at −54 kJ/mol (ΔGATP_-54_), or following the addition of complex I substrates (octanoyl-carnitine [Oct-Carn], glutamate [Glut]), complex II substrates (succinate [Succ]), the respiratory uncoupler FCCP [FC], or the complex I inhibitor rotenone. Note different y-axis ranges in panels **C** versus **D**. Data are presented as mean ± SD. SAC = SACLAC; LCL = LCL-805; VEN = venetoclax. Concentrations in brackets are µM. *p<0.05, **p<0.01, ***p<0.005, ****p<0.0001, ns = not significant.

We next investigated the effect of AC and BCL-2 co-targeting on mitochondrial respiration using a comprehensive mitochondrial diagnostic workflow to assess contributions to the combined cytotoxic mechanism (40, 43). This approach leveraged a modified version of the creatine kinase clamp to titrate extramitochondrial ATP/ADP ratios and mimic physiologically relevant ATP free energies (ΔG_ATP_). Following AC inhibition or venetoclax treatment, cells were permeabilized with digitonin and oxygen consumption was measured after respiratory stimulation by clamping ΔG_ATP_ at −54 kJ/mol, supplementing cells with mitochondrial electron transport complex-specific substrates, or treating cells with the respiratory uncoupler FCCP. In MM-6 cells, treatment with two AC inhibitors, SACLAC or LCL-805, partially antagonized respiration irrespective of the stimulant (**Fig. 4c**). Venetoclax alone was sufficient to significantly reduce respiration, which was enhanced by combining venetoclax with either SACLAC or LCL-805 (**Fig. 4c**). Though SACLAC, LCL-805, venetoclax, SACLAC + venetoclax, and LCL-805 + venetoclax inhibited respiration in MV4-11 VEN-R cells, the effects were modest due to the poor respiratory capacity of this cell line. In contrast to MM-6, the stimulants failed to substantially increase mitochondrial respiration in MV4-11 VEN-R cells (**Fig. 4d**). These findings are consistent with our previous report that blocking mitochondrial respiration is not sufficient to induce AML cell death (44). These findings highlight i) stark differences in mitochondrial respiratory capacity between MM-6 and MV4-11 VEN-R cells, ii) that venetoclax alone was sufficient to reduce mitochondrial respiration to near baseline in MM-6 cells and respiration was very low in MV4-11 VEN-R though both cell lines remained viable, and iii) that the addition of two independent AC inhibitors resulted in additional loss of respiratory capacity beyond that which was observed with venetoclax alone.

### AC inhibition enhanced the efficacy of clinically relevant venetoclax and cytarabine combination

Venetoclax is FDA-approved in combination with low-dose cytarabine for AML patients ineligible for intensive chemotherapy (45) and the efficacy of this combination is enhanced by exogenous ceramide supplementation in various AML models (19). To explore how AC inhibition augments relevant AML therapeutics, we expanded our study to include drug combinations with cytarabine. MM-6 and MV4-11 VEN-R cells were treated with the three-drug combination of SACLAC + venetoclax + cytarabine, each two-drug permutation, and single agent or vehicle controls. As single agents, SACLAC, venetoclax, and cytarabine modestly reduced cell viability and increased the fraction of annexin V-positive cells. The two-drug combinations of SACLAC + venetoclax and SACLAC + cytarabine exhibited similar efficacy to venetoclax + cytarabine. Importantly, the triple-drug combinations resulted in significantly decreased cell viability and increased cytotoxicity compared to two-drug combinations or single-agent controls (**Fig. 5a-b**). These data show that AC inhibition enhanced the cytotoxicity of venetoclax and cytarabine as single agents and significantly augmented the cytotoxicity of the venetoclax + cytarabine combination.

**Fig 5:**
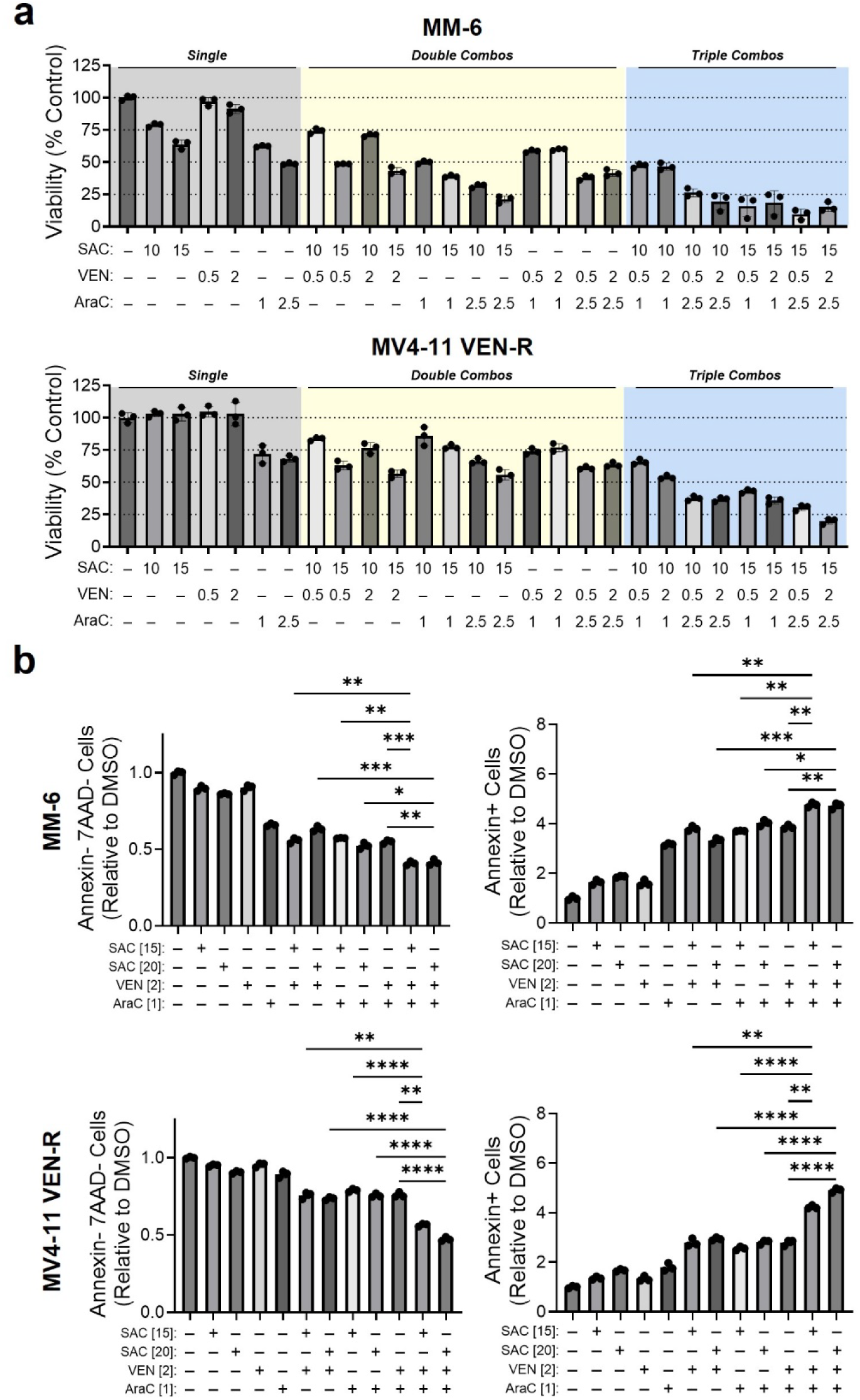
Acid ceramidase inhibition augmented the cytotoxicity of venetoclax and cytarabine. **A** MM-6 and MV4-11 VEN-R cells were treated with DMSO, SACLAC, venetoclax, cytarabine, doublet combinations, and triplet combinations at the indicated concentrations for 48 h then MTS cell viability was assessed. **B** Flow cytometry annexin V profiling of MM-6 and MV4-11 VEN-R cells treated with DMSO, SACLAC, venetoclax, cytarabine, doublet combinations, and triplet combinations at the indicated concentrations for 48 h. Significance was assessed by Welch’s ANOVA with Dunnet’s multiple comparisons test. Data are presented as mean ± SD. SAC = SACLAC; VEN = venetoclax; Ara-C = cytarabine. Concentrations in brackets are µM. *p<0.05, **p<0.01, ***p<0.005, ****p<0.0001, ns = not significant.

### Co-targeting AC and BCL-2 increased cytotoxicity in primary AML patient samples

We extended our efficacy studies beyond cell lines and performed *ex vivo* drug screening of SACLAC, venetoclax, and cytarabine across a cohort of primary AML patient and healthy control samples. Our AML cohort consisted of peripheral blood mononuclear cells (PBMCs) and bone marrow (BM)-derived cells from **71** patients. We also screened 7 PBMC and 2 BM samples from healthy individuals. We previously reported patient demographic and clinical information for this cohort (23). We first determined IC_50_ values for SACLAC, venetoclax, or cytarabine. SACLAC was significantly more toxic toward samples derived from AML patients versus healthy control samples (**Fig. 6a**). Venetoclax and cytarabine IC_50_ values were highly heterogenous and the mean IC_50_ values were lower in AML samples versus healthy controls but not statistically significant (**Fig. 6b-c**). We next tested the SACLAC + venetoclax combination across the AML specimens and healthy control samples. In line with our cell line data, SACLAC + venetoclax treatment exhibited greater efficacy than SACLAC or venetoclax alone. The AC inhibitors SACLAC and LCL-805 also reduced colony formation potential in primary AML patient samples (**Fig. S3d-e**). Importantly, SACLAC + venetoclax was significantly more toxic to AML specimens than healthy control samples (**Fig. 6d**). We also compared the SACLAC + venetoclax combination versus the clinically utilized venetoclax + cytarabine combination and found that they comparably reduced cell viability in patient samples (**Fig. 6e**). Bliss synergy analysis revealed enhanced synergy scores of the SACLAC + venetoclax combination versus venetoclax + cytarabine for most patient samples (**Fig. 6f**). Together, these findings demonstrate promising efficacy of SACLAC + venetoclax at reducing primary AML patient sample cell viability.

**Fig. 6:**
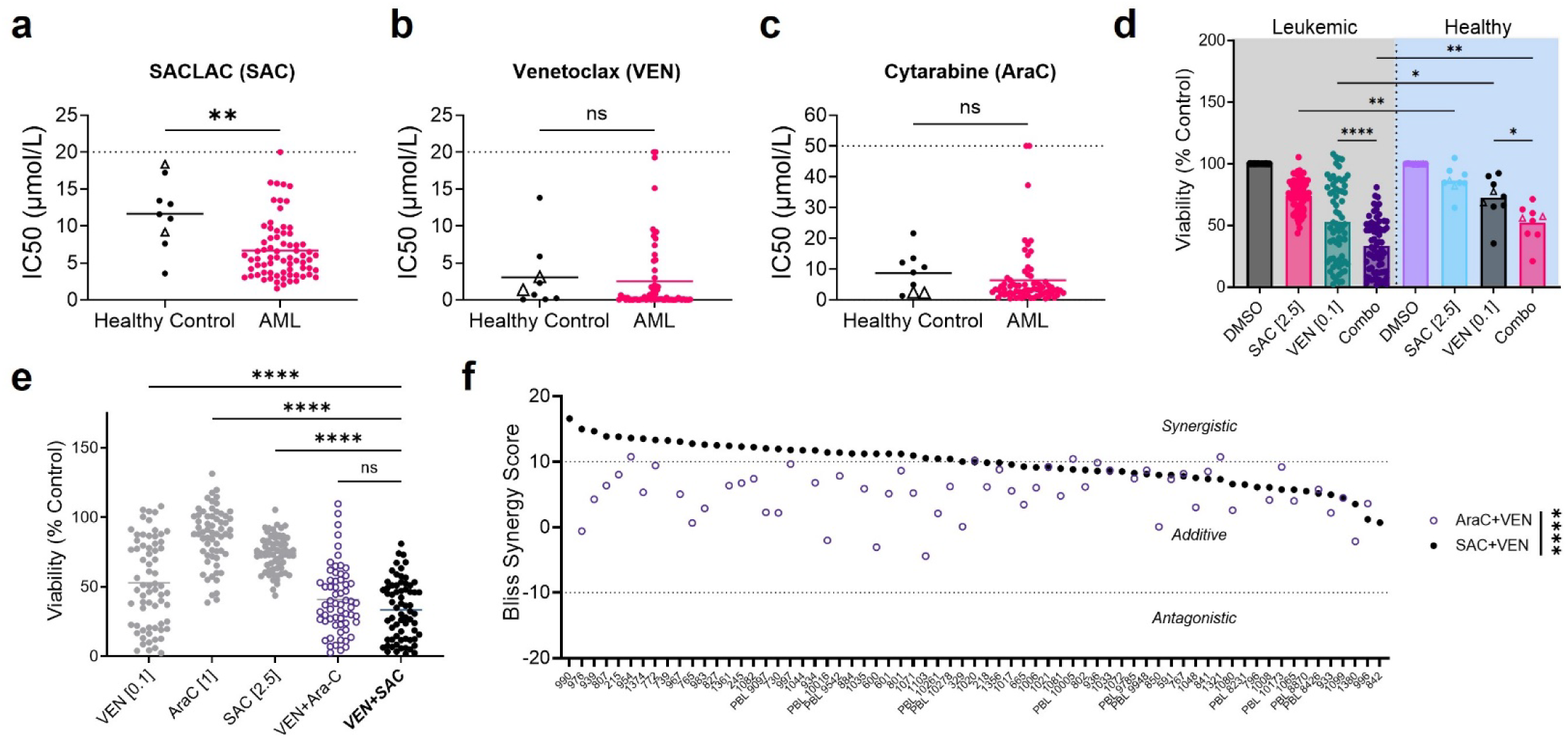
SACLAC enhanced venetoclax cytotoxicity in primary acute myeloid leukemia patient samples. **A-C** Primary patient samples (n=71) and healthy PBMC (n=7) or BM (n=2) control samples were treated with increasing concentrations of SACLAC, venetoclax, or cytarabine for 48 h after which cell viability was assessed by CellTiterGlo. IC_50_ values were determined for each drug using the GraphPad Prism “log(inhibitor) vs. normalized response -- Variable slope” function. Dotted line represents the highest concentration tested and samples on the dotted line did not reach IC_50_ using tested concentrations. Significance was assessed with the Mann-Whitney test. **D** Primary patient samples and healthy PBMC or BM control samples were treated with DMSO, SACLAC, venetoclax, or the combination for 48 h after which cell viability was assessed by CellTiterGlo. Significance was assessed by unpaired t-test with Welch’s correction. **E, F** Primary patient samples were treated with DMSO, SACLAC, cytarabine, venetoclax, or doublet combinations for 48 h after which cell viability was assessed by CellTiterGlo. Significance was assessed by unpaired t-test with Welch’s correction (**E**) or Wilcoxon matched-pairs signed rank test (**F**). Bliss synergy scores were calculated using SynergyFinder 2.0. Bliss scores greater than 10 were considered synergistic, between −10 and 10 were considered additive, and less than −10 were considered antagonistic. Healthy control bone marrow samples were denoted with an open triangle and all other healthy control samples were PBMC-derived. Data are presented as mean ± SD. SAC = SACLAC; VEN = venetoclax; Ara-C = cytarabine. Concentrations in brackets are µM. *p<0.05, **p<0.01, ***p<0.005, ****p<0.0001, ns = not significant.

## Discussion

The adoption of venetoclax-containing regimens improved the treatment landscape for AML patients unfit for intensive combination chemotherapy (45). Though initially effective, 30-40% of patients are refractory to treatment and most patients develop resistance (25). Thus, understanding regulators of venetoclax efficacy in AML is of high importance (10, 25). In this work, we demonstrated the efficacy of co-targeting AC and BCL-2 across AML cell lines and primary patient samples. Pharmacologic AC inhibition enhanced both the efficacy of venetoclax and venetoclax + cytarabine in venetoclax-resistant AML cell lines. Central to the cytotoxic mechanism were dramatic increases of intracellular ceramides and induction of a cytotoxic ISR leading to NOXA accumulation, mitochondrial depolarization, and caspase-dependent cell death.

Metabolic rewiring is an emergent AML hallmark, and dysregulated sphingolipid metabolism is becoming recognized as an important feature (20, 46). Ceramides are tumor-suppressive sphingolipids that play a key cytotoxic role in AML-relevant therapeutics. For example, the anthracycline daunorubicin is used extensively for AML induction therapy and the intracellular ceramide it generates contributes to its cytotoxic mechanism (15, 16). This “ceramide-killing effect” is not limited to chemotherapeutic drugs. Elevated ceramide levels, induced by antagonizing ceramide glycosylation or sphingosine kinases, also improve BH3 mimetic efficacy across various human leukemias (47). Despite this, few studies have targeted ceramide metabolism to augment venetoclax efficacy. We previously showed that venetoclax treatment in combination with exogenous C6-ceramide nanoliposomes (CNL) enhanced intracellular ceramide accumulation to support the combined cytotoxic mechanism (19). We also previously identified the ceramide-catabolizing lipid hydrolase, AC, as a mediator of BH3 mimetic ABT-737 efficacy in AML (17). However, the link between AC targeting and venetoclax in AML remained unexplored until now.

Ceramides are hydrolyzed by ceramidases into sphingosine, which serves as the substrate for sphingosine kinase-mediated phosphorylation to S1P. Conversely, S1P is dephosphorylated by S1P phosphatases to generate sphingosine, which is acylated by ceramide synthases to produce ceramides. In the original cell-intrinsic rheostat model, increased ceramides and decreased S1P are considered pro-death while decreased ceramide and increased S1P are considered pro-survival (48). Although the rheostat model has been updated to reflect new research (e.g., different ceramide species exert distinct functions; inside-out S1P paracrine signaling) (48), therapeutic strategies aimed at increasing total intracellular ceramides or decreasing S1P production are promising strategies for AML as single agents or in combination with venetoclax (22, 28, 49). CNL enhanced venetoclax and cytarabine efficacy in AML (19) and has now entered a phase I clinical trial for relapsed/refractory AML (NCT04716452). Others demonstrated the utility of inhibiting sphingosine kinase to augment venetoclax killing in AML (28). We extend this work by 1) targeting AC, which is upstream of sphingosine kinase in the salvage pathway, 2) evaluating AC and BCL-2 co-targeting in venetoclax-resistant AML, 3) linking synergy directly to the ISR, 4) extending studies to a large cohort of primary patient samples, and 5) assessing AC inhibition in the context of the FDA-approved venetoclax + cytarabine combination. Collectively, these studies demonstrate that inhibiting flux through the sphingolipid salvage pathway or supplementing with exogenous ceramide are promising approaches to improve venetoclax cytotoxicity in venetoclax-sensitive and -resistant AML cells. Whereas we evaluated sphingolipid changes following SACLAC and venetoclax treatment in these cells, the complete set of sphingolipid metabolic alterations and flux changes in primary and acquired venetoclax resistance is unknown and an open area of study.

ISR signaling is emerging as a crucial regulator of BH3 mimetic efficacy. Indeed, ISR activation was sufficient to enhance venetoclax toxicity in venetoclax-resistant AML cell lines (33). Moreover, the sphingosine kinase inhibitor MP-A08 sensitized AML cells to venetoclax-induced cytotoxicity via a PKR-mediated cytotoxic ISR (28). We also identified the ISR as a mediator of efficacy between AC and BCL-2 co-targeting through rescue experiments with the ISR inhibitor ISRIB. Our data demonstrate that PKR inhibition blunted ATF4 and NOXA induction following AC inhibition. Ceramides have previously been shown to induce PKR activation and blunt protein synthesis (50). In addition to PKR, our findings highlight an additional link between AC inhibition, ceramides, and ISR signaling through GCN2 as we found that GCN2 inhibition with GCN2-IN-1 also blocked ATF4 and NOXA induction. Indeed, a past study evaluated ceramide binding partners and predicted binding between ceramide and GCN1 (51), a critical regulator of GCN2 activity (52). In support of this GCN2 model, we observed that SACLAC treatment led to increased protein levels of GCN1 and GIGYF2, which serve to activate GCN2 and respond to ribosomal stress (37). Collectively, these findings reveal a novel relationship between ceramide-induced ribosomal stress and the ISR.

Venetoclax resistance is associated with upregulation of anti-apoptotic proteins such as MCL-1 and BCL-xL (10, 25). Unfortunately, BCL-xL inhibition in leukemia is limited by on-target dose-limiting thrombocytopenia (53). Strategies targeting MCL-1 are actively being investigated preclinically and clinically though development may be hindered by cardiac toxicities (54). ISR activation is linked to both MCL-1-dependent and independent mechanisms of improving venetoclax killing. For example, sphingosine kinase inhibition enhanced venetoclax killing through a PKR/ATF4/NOXA/MCL-1 axis (28). In a separate study, targeting mitochondrial translation with antibiotics enhanced venetoclax killing by inhibiting mitochondrial respiration and causing ISR-mediated glycolytic impairment independent of MCL-1 or BCL-xL protein expression changes (33). In our model, MCL-1 or BCL-xL protein levels did not decrease in response to AC inhibition and/or venetoclax treatment in MM-6 and MV4-11 VEN-R cells. Instead, cell killing from the combination co-occurred with ISR-mediated upregulation of the endogenous MCL-1 inhibitor, NOXA, and loss of mitochondrial membrane potential. The observation that MM-6 and MV4-11 VEN-R cells survive acute venetoclax exposure despite reduced mitochondrial respiration upon single-agent venetoclax treatment suggests that AC-mediated enhancement of venetoclax efficacy may engage additional mechanisms beyond further inhibition of mitochondrial respiration. The findings here are consistent with our recent report that venetoclax-resistant AML cells hydrolyze ATP to maintain a mitochondrial membrane potential and resist venetoclax-induced cell death (44). Instead of targeting respiration, targeting mitochondrial membrane potential was key to inducing AML cell death (44). This may suggest that AC inhibition enhances venetoclax cytotoxicity by preventing cells from hydrolyzing ATP to maintain a mitochondrial membrane potential and resist BCL-2 targeting. Collectively, the cytotoxic mechanism of AC targeting and BCL-2 inhibition likely incorporates both ISR-mediated NOXA protein upregulation and more complex effects on mitochondrial polarization.

Intensive chemotherapy combining cytarabine and an anthracycline remains the backbone of AML induction therapy (45). We previously reported that cytarabine and daunorubicin selection pressure promotes a pro-survival sphingolipid composition that reduces intracellular ceramide levels in part through increased AC activity and protein expression (14). These adaptations highlight a potentially exploitable vulnerability to improve chemotherapy efficacy in AML. Indeed, our efficacy studies demonstrated that AC inhibition synergized with cytarabine alone. The polychemotherapy consisting of SACLAC + venetoclax + cytarabine was more efficacious than either doublet combination or single agent control. The pro-survival changes in sphingolipid metabolism in chemotherapy-resistant AML also include other pathways that decrease intracellular ceramide levels (14), thus warranting additional studies that evaluate other sphingolipid modulating drugs (e.g., glucosylceramide synthase inhibitors, sphingosine kinase inhibitors, ceramide kinase inhibitors) in the context of venetoclax-containing regimens.

In summary, we identified AC as a regulator of venetoclax efficacy in venetoclax-resistant AML. Pharmacologic inhibition of AC was sufficient to enhance venetoclax-mediated killing via a cytotoxic ISR and mitochondrial impairment. AC inhibition also increased AML sensitivity to cytarabine and the venetoclax + cytarabine combination. These findings support the continued development of sphingolipid inhibitors to augment approved AML drugs.

## Acknowledgments

The authors would like to acknowledge and thank the patients and their families who supported our studies. We thank Gemma Fabrias for providing SACLAC and Galina Diakova, the UVA Partners in Discovery Team, the UVA Office of Clinical Research Non-Treatment Research Operations, and the UVA Biorepository and Tissue Research Facility in the consenting of patients, specimen procurement, specimen processing, data abstraction, and providing access to molecular and clinical data (IRB-HSR #18445). This work was supported by the Veteran’s Administration (VA Merit Review, I BX001792 [to CEC]) and a Research Career Scientist Award, IK6BX004603 [to CEC]) and the National Institutes of Health (NIH) award numbers P01 CA171983 (to TPL and CEC), Cancer Center Support Grant P30 CA044579 (to TPL), R01 AI139072 (to CEC), F31 CA271809 (to JU), and F99 CA284252 (to JU) This work was also supported by the UVA Robert R. Wagner Fellowship (to JU). The content is solely the responsibility of the authors and does not necessarily represent the official views of the National Institutes of Health, Department of Veterans Affairs, or the United States Government. This work is dedicated in loving memory to the late sphingolipid pioneer Dr. Mark Kester.

## Contributions

JU, SFT, DJF, and TPL conceptualized the study design. JU, SFT, JJPS, MT, TMD, GDCV, MMM, JTH, RTA, URG, AS, BBP, ISL, BR, KFW, and TEF were responsible for experimental work, data collection, and data analysis. JU, KFW, and BBP performed proteomic, statistical, and bioinformatic analyses. KAJ, FGB, MCC, and DFC provided scientific resources and subject matter expertise. DJF and TPL provided project oversight. JU, CEC, and TPL were responsible for funding acquisition. JU drafted the original manuscript draft. All co-authors provided edits, reviewed, and approved the final manuscript.

## Data Availability Statement

Data needed to evaluate conclusions from this work are present in the paper. Molecular and clinical characteristics from the primary AML patient samples were previously reported (23). All proteomics data will be deposited in ProteomeXchange six months after publication. Summary data for pathway analysis is available in Table S1. Other datasets generated during and/or analyzed during the current study are available from the corresponding author on reasonable request.

---

**Fig. S1:**
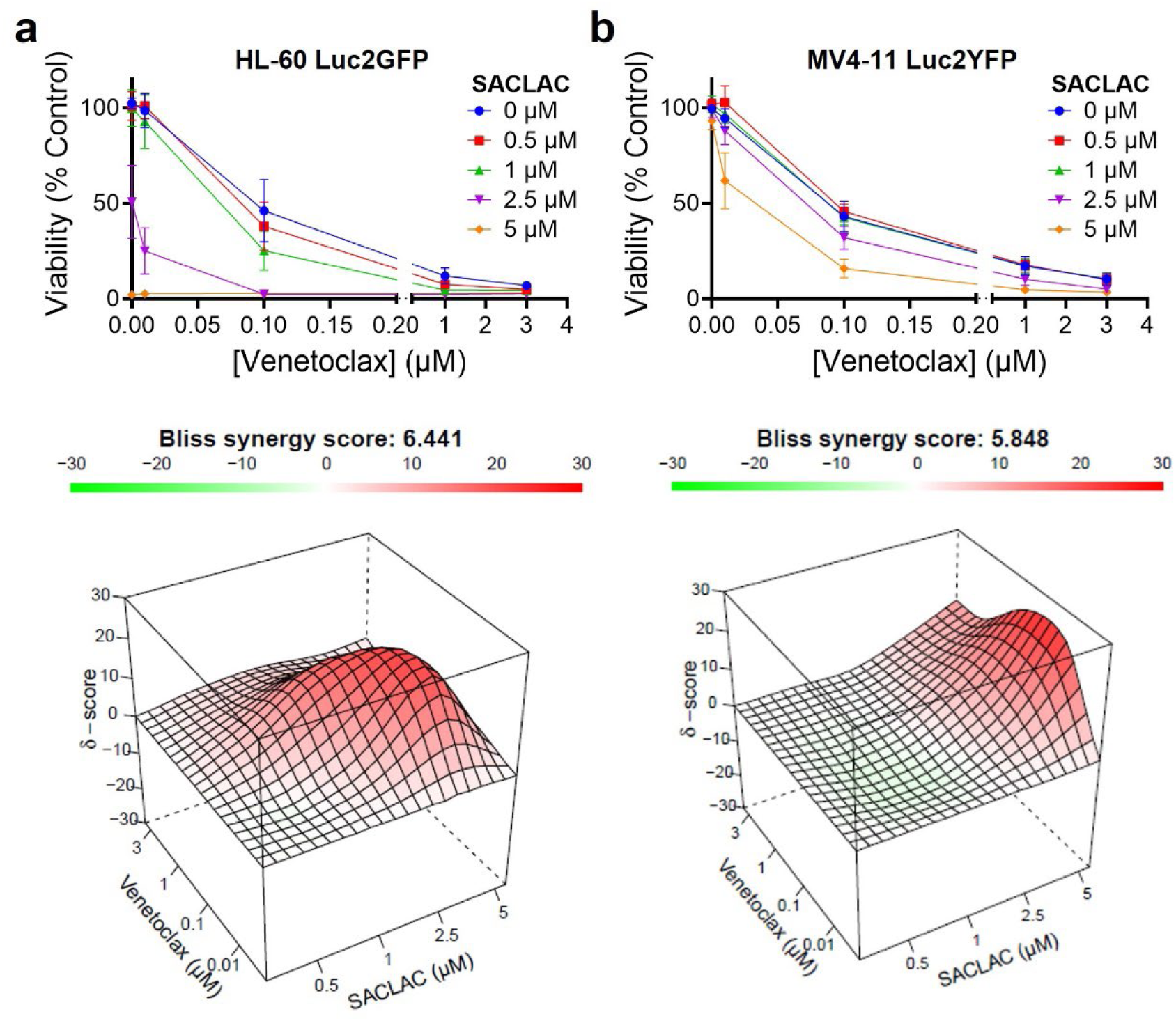
Pharmacologic inhibition of acid ceramidase enhanced venetoclax cytotoxicity in venetoclax-sensitive acute myeloid leukemia cell lines. HL-60 Luc2GFP **(A)** and MV4-11 Luc2YFP **(B)** cells were treated with increasing concentrations of SACLAC, venetoclax, or the combination for 48 h and cell viability was assessed by MTS. Bliss synergy scores were calculated using SynergyFinder 3.0. Bliss scores greater than 10 were considered synergistic, between −10 and 10 were considered additive, and less than −10 were considered antagonistic. Data are presented as mean ± SD.

**Fig. S2:**
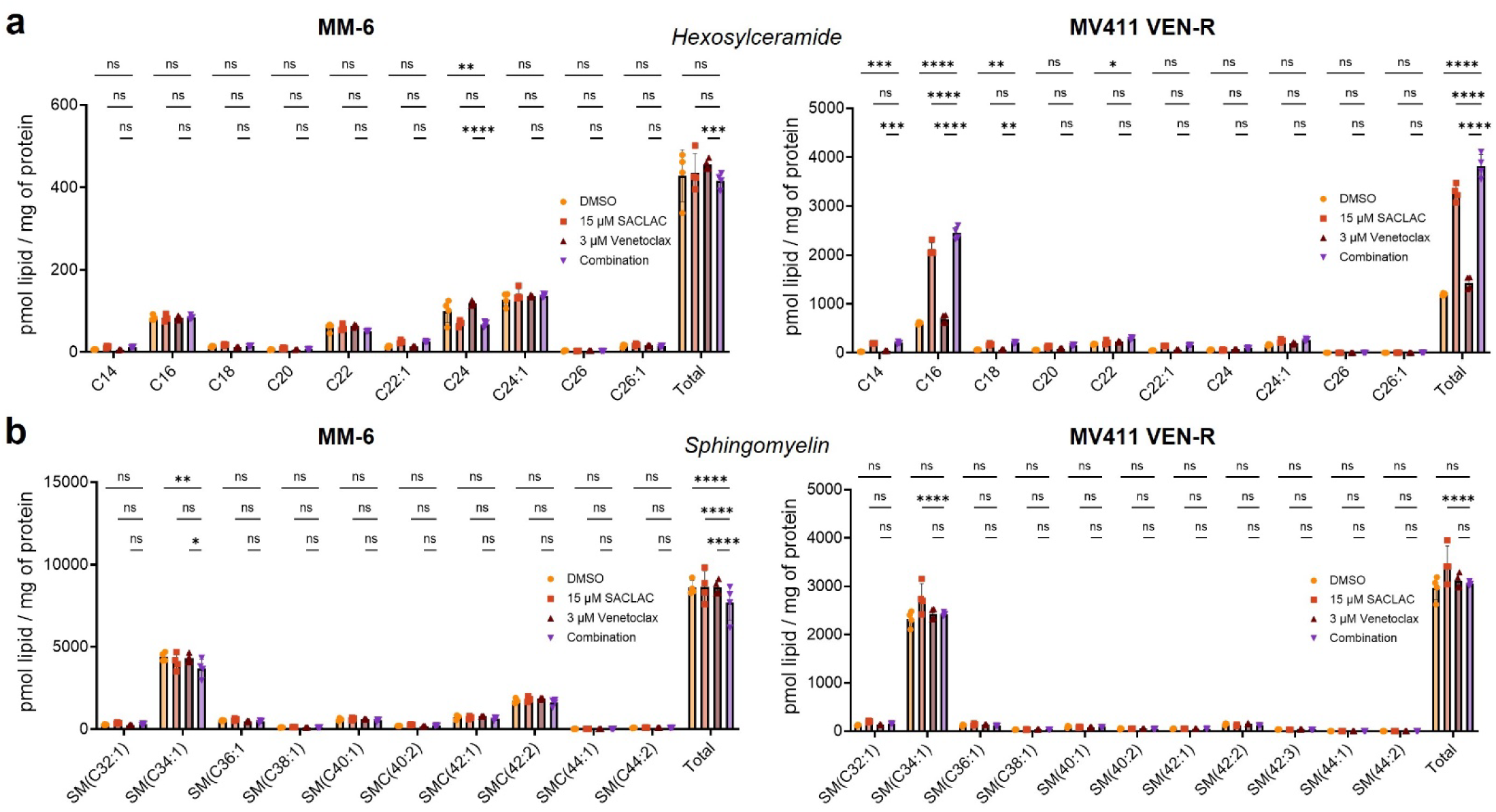
Effect of co-targeting acid ceramidase and BCL-2 on hexosylceramide and sphingomyelin levels. MM-6 and MV4-11 VEN-R cells were treated with DMSO, SACLAC, venetoclax, or the combination for 24 h, and hexosylceramide **(A)** or sphingomyelin **(B)** levels were measured by liquid chromatography-mass spectrometry. Significance was assessed by two-way ANOVA with Tukey’s multiple comparisons test. *p<0.05, **p<0.01, ***p<0.005, ****p<0.0001, ns = not significant.

**Fig. S3:**
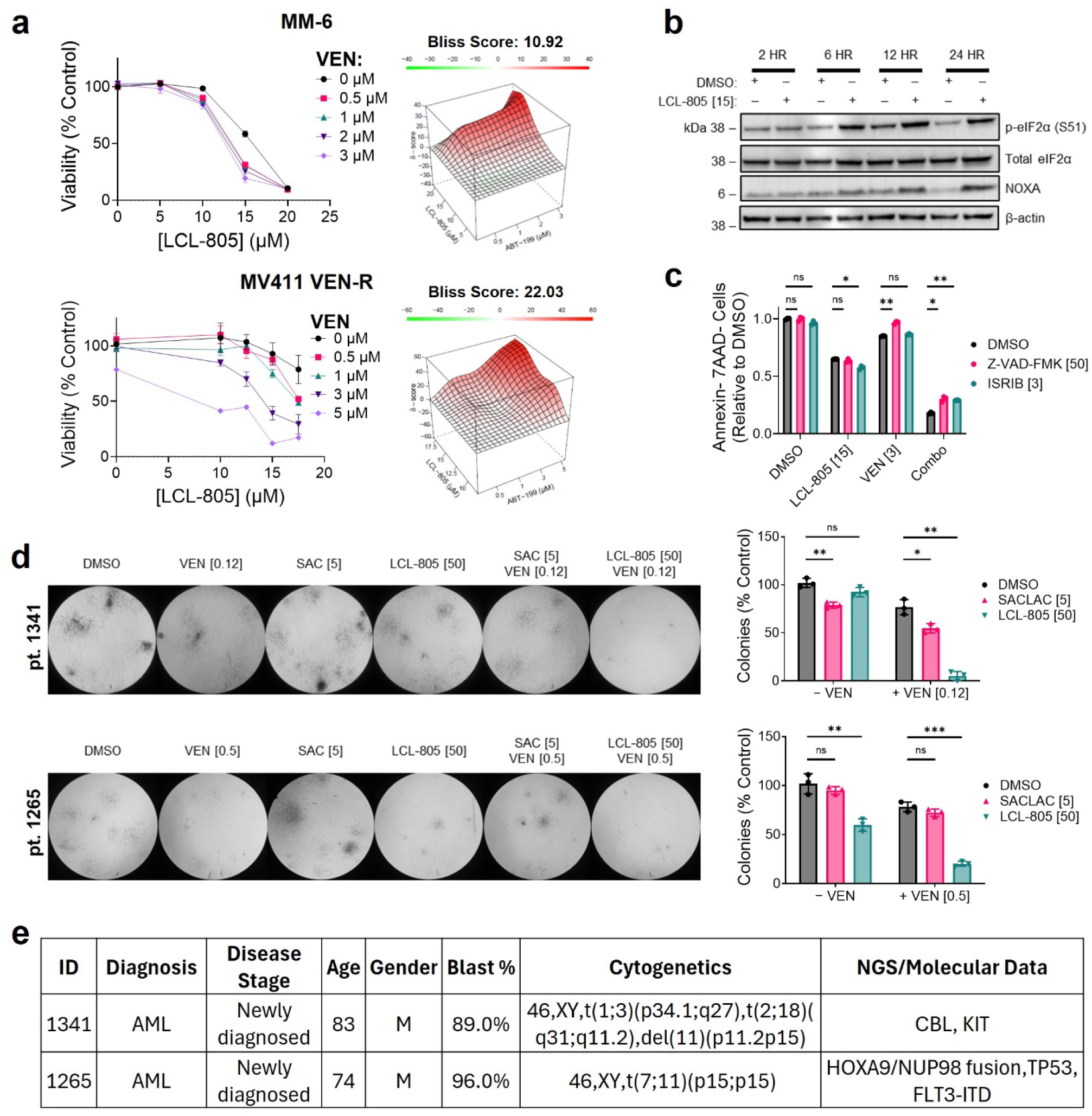
LCL-805 enhanced venetoclax-mediated cytotoxicity via caspase activation and a cytotoxic integrated stress response. **A** MM-6 and MV4-11 VEN-R cells were treated with DMSO, LCL-805, venetoclax, or the combination for 24 h or 72 h, respectively, and cell viability was assessed by MTS. Bliss synergy scores were calculated using SynergyFinder 2.0. Bliss scores greater than 10 were considered synergistic, between −10 and 10 were considered additive, and less than −10 were considered antagonistic. **B** Immunoblotting of MM-6 cells treated with DMSO or LCL-805 at the indicated concentrations and times. β-actin expression served as the loading control for immunoblots. **C** Flow cytometry profiling of annexin V/7-AAD negative cells pretreated MM-6 cells were pretreated with DMSO, Z-VAD-FMK, or ISRIB for 2 h followed by DMSO, LCL-805, venetoclax, or combination treatment for 24 h. Significance was assessed with unpaired Welch’s t-test with Holm-Šídák correction. **D** Clonogenic capacity of primary AML patient samples treated with DMSO, SACLAC, LCL-805, venetoclax, SACLAC + venetoclax, or LCL-805 + venetoclax at the indicated concentrations. Significance was assessed with unpaired Welch’s t-test with Holm-Šídák correction. **E** Clinical data for patient samples 1265 and 1341. Data are presented as mean ± SD. SAC = SACLAC; ABT-199 = venetoclax (VEN). Concentrations in brackets are µM. *p<0.05, **p<0.01, ****p<0.0001, ns = not significant.

