## Supplemental Methods for "Acid ceramidase inhibition enhances BCL-2 targeting in venetoclax-resistant acute myeloid leukemia via a cytotoxic integrated stress response"

**Running Title:** Co-Targeting Acid Ceramidase and BCL-2 in AML


**Author List and Affiliations:**

Johnson Ung[1], Su-Fern Tan[2,3], Jeremy J.P. Shaw[2,3], Maansi Taori[2,3], Tess M. Deddens[2,3], Giovana da Costa Venancio[2,3], McLane M. Montgomery[4,5], James T. Hagen[4,5], Raphael T. Aruleba[6], Upendar R. Golla[7,8], Arati Sharma[8,9], B. Bishal Paudel[2,3,10], Irene Sung-Ah Lee[2,3], Bhavishya Ramamoorthy[2,3], Kevin A. Janes[10,11], Francine Garrett-Bakelman[2,3,11], Myles C. Cabot[3], Kelsey H. Fisher-Wellman[6], Todd E. Fox[12], David F. Claxton[7,8], Charles E. Chalfant[2,3], David J. Feith[2,3], Thomas P. Loughran Jr.[2,3]

[1]Department of Microbiology, Immunology, and Cancer Biology, University of Virginia School of Medicine, Charlottesville, VA, USA
[2]Department of Medicine, Division of Hematology & Oncology, University of Virginia School of Medicine, Charlottesville, VA, USA
[3]University of Virginia Comprehensive Cancer Center, Charlottesville, VA, USA
[4]Department of Physiology, Brody School of Medicine, East Carolina University, Greenville, NC, USA
[5]East Carolina Diabetes and Obesity Institute, East Carolina University, Greenville, NC, USA
[6]Department of Cancer Biology, Atrium Health Wake Forest Baptist Comprehensive Cancer, Wake Forest University School of Medicine, Winston-Salem, NC, USA
[7]Division of Hematology and Oncology, Department of Medicine, Pennsylvania State University College of Medicine, Hershey, PA, USA
[8]Penn State Cancer Institute, Pennsylvania State University College of Medicine, Hershey, PA, USA
[9]Department of Molecular and Precision Medicine, Pennsylvania State University College of Medicine, Hershey, PA, USA
[10]Department of Biomedical Engineering, University of Virginia, Charlottesville, VA, USA
[11]Department of Biochemistry and Molecular Genetics, University of Virginia School of Medicine, Charlottesville, VA, USA
[12]Department of Pharmacology, University of Virginia School of Medicine, Charlottesville, VA, USA

**Corresponding Author:**
Thomas P. Loughran, Jr., M.D.
Department of Medicine, Division of Hematology & Oncology
University of Virginia School of Medicine, Charlottesville, VA 22908

### Supplemental Materials and Methods:

#### Reagents

Inhibitors: SACLAC was kindly provided by Gemma Fabrias and purchased from AmBeed, Arlington Heights, IL; venetoclax (Selleck Chemicals, Houston, TX, USA, # S8048); cytarabine (Selleck Chemicals, # S1648); Z-VAD-FMK (Selleck Chemicals, #S7023); ISRIB (Sigma, St. Louis, MO, USA, #SML0843); vincristine (Cayman, Ann Arbor, MI, USA, # 11764)

Antibodies from Cell Signaling Technologies (Danvers, MA, USA) include: PARP (#9542); Caspase-9 (#9508); Caspase-3 (#9662); NOXA (#14766); MCL-1 (#5453); BCL-2 (#4223); Bcl-xL (#2764); p-eIF2α Ser51 (#3597); eIF2α (#5324); ATF4 (#11815); β-actin (#3700); anti-rabbit IgG, HRP-linked (#7074); and anti-mouse IgG, HRP-linked (#7076). Anti-acid ceramidase (#612302) was acquired from BD Transduction Laboratories (Franklin Lakes, NJ, USA).

#### Patient samples

Informed consent for sample collection and testing was obtained in accordance with approved protocols via the institutional review boards of the Penn State Hershey Medical Center and the University of Virginia School of Medicine and in accordance with the Declaration of Helsinki. Healthy control peripheral blood mononuclear cells (PBMCs) and bone marrow (BM) samples were obtained from AllCells (Alameda, CA, USA), Virginia Blood Services, Zen-Bio (Durham, NC, USA), and Inova Blood Donor Services (Sterling, VA, USA). Cells were isolated using the Ficoll-Paque gradient separation method and cryopreserved. Cryopreserved normal and AML patient PBMC samples were thawed and resuspended in RPMI supplemented with 15% FBS.

#### Colony Formation Assays

The clonogenicity of primary AML patient cells was tested in the presence of the indicated drug treatments and doses by seeding 1,000 cells per well in Human Methylcellulose Complete Media (R&D Systems, Minneapolis, MN, USA) in a 12-well plate. The plates were kept in cell culture incubator (37oC with 5% CO2) for 10-14 days and the resultant blast colonies (>20 cells/colony) were counted under an Olympus CKX31 inverted light microscope (Olympus Corporation, Center Valley, PA, USA) and imaged using a 4x objective. The data was represented as percentage colonies relative to solvent (DMSO) control.

**Proteomics analysis**

Cell pellets were lysed in urea lysis buffer (8 M urea in 40 mM Tris, 30 mM NaCl, 1mM CaCl2, 1 x cOmplete ULTRA mini EDTA-free protease inhibitor tablet; pH=8.0). The samples were subjected to two freeze-thaw cycles, and sonicated with a probe sonicator in three 5 s bursts (Q Sonica #CL-188; amplitude of 30). Samples were centrifuged at 10,000 x g for 10 min at 4°C. Protein concentration was determined by BCA. Equal amounts of protein were reduced with 5 mM DTT at 37°C for 30 min, and then alkylated with 15 mM iodoacetamide for 30 min in the dark. Unreacted iodoacetamide was quenched with DTT [15 mM]. Reduction and alkylation reaction were carried out at room temperature. Initial digestion was performed with Lys C (1:100 w:w) for 4 h at 32°C. Following dilution to 1.5 M urea with 40 mM Tris (pH=8.0), 30 mM NaCl, 1 mM CaCl2, samples were digested overnight with sequencing grade trypsin (50:1 w/w) at 32°C. Samples were acidified to 0.5% TFA and then centrifuged at 4,000 x g for 10 min at 4°C. Supernatant containing soluble peptides was desalted and then eluate was frozen and subjected to speedvac vacuum concentration. Final peptides were resuspended in 0.1% formic acid, quantified (Thermo Fisher Scientific; 23275), and diluted to a final concentration of 0.25 µg/µL. Samples were subjected to nanoLC-MS/MS analysis using an UltiMate 3000 RSLCnano system (Thermo Fisher Scientific) coupled to a Q Exactive Plus Hybrid Quadrupole-Orbitrap mass spectrometer (Thermo Fisher Scientific) via a nanoelectrospray ionization source. For each injection, 4µL (1µg) of sample was first trapped on an Acclaim PepMap 100 20mm × 0.075mm trapping column (Thermo Fisher Scientific; 164535; 5µL/min at 98/2 v/v water/acetonitrile with 0.1% formic acid). Analytical separation was performed over a 95 min gradient (flow rate of 250nL/min) of 4-25% acetonitrile using a 2 µm EASY-Spray PepMap RSLC C18 75µm × 250mm column (Thermo Fisher Scientific; ES802A) with a column temperature of 45ºC. MS1 was performed at 70,000 resolution, with an AGC target of 3x106 ions and a maximum injection time (IT) of 100ms. MS2 spectra were collected by data-dependent acquisition (DDA) of the top 15 most abundant precursor ions with a charge greater than 1 per MS1 scan, with dynamic exclusion enabled for 20s. Precursor ions isolation window was 1.5m/z and normalized collision energy was 27. MS2 scans were performed at 17,500 resolution, maximum IT of 50ms, and AGC target of 1x105 ions. Proteome Discoverer 2.2 (PDv2.2) was used for raw data analysis, with default search parameters including oxidation (15.995 Da on M) as a variable modification and carbamidomethyl (57.021 Da on C) as a fixed modification. Data were searched against the Uniprot Homo Sapiens reference proteome (Proteome ID: UP000005640). PSMs were filtered to a 1% FDR and grouped to unique peptides while maintaining a 1% FDR at the peptide level. Peptides were grouped to proteins using the rules of strict parsimony and proteins were filtered to 1% FDR. Peptide quantification was done using the MS1 precursor intensity. Imputation was performed via low abundance resampling. All proteomics samples were normalized to total protein abundance, and the protein tab in the PDv2.2 results was exported as a tab delimited .txt. file and analyzed. Protein abundance was converted to the log2 space. For pairwise comparisons, cell mean, standard deviation, p-value (p; two-tailed Student's t-test, assuming equal variance), and adjusted p-value (Benjamini Hochberg FDR correction) were calculated.

**Mitochondrial profiling**

Respirometry measurements were conducted using an Oroboros Oxgraph-2k (O2k; Oroboros Instruments, Innsbruck, Austria) in either a 0.5 or 1.0mL reaction volume at 37°C. Data was normalized to viable cell count using trypan blue exclusion. For each assay, cells were suspended in either 0.5 mL or 1 mL of Respiration Buffer supplemented with creatine (105 mM MES potassium salt, 30 mM KCl, 8 mM NaCl, 1 mM EGTA, 10 mM $KH_2PO_4$, 5mM $MgCl_2$, 0.25% BSA, 5 mM creatine monohydrate, pH 7.2). Cells were then permeabilized with digitonin (0.015mg/mL). The OxPhos system was stimulated using a modified version of the creatine kinase clamp technique. The free energy of ATP hydrolysis (ΔGATP) was calculated using the equilibrium constant for the CK reaction (K'CK) and is based upon the addition of known concentrations of creatine (CR 5mM), phosphocreatine (PCR 1mM), and ATP (10mM) in the presence of excess amounts of CK (20U/mL). Calculation of ΔGATP was done as previously described [107]. Additional substrates/inhibitors added during the assay were as follows: octanoyl-carnitine (Oct-Carn; 0.2 mM), succinate (Succ or S; 5 mM), glutamate (Glut; 5 mM), rotenone (Rot; 0.5 μM), and FCCP titration (FC, 0.5-2 μM).

**Cell transduction protocol**

To generate HL-60 LUC2-eGFP cells, $1x10^6$ cells in 6-well plates were transduced with lentivirus containing the pFU-LUC2-eGFP plasmid. 8 µg/ml polybrene was added to the plate and centrifuged at 1100 rpm for 1 hr. EGFP-positive cells were confirmed at 72 h post-transduction and flow-sorted for positive GFP expression.

**Enrichment analysis using STRINGDb**

Differentially upregulated (n = 249) and downregulated (n = 190) proteins were analyzed for enrichment using STRING database website (PMID: 30476243) under Search>Multiple proteins. Only the proteins that mapped with the STRING database were kept, resulting in 247 upregulated, and 186 downregulated proteins. The default settings were used for analyses with minimum required interactions score set to medium confidence (0.4).
